## Supplementary Figures and Tables for "A practical DNA data storage using expanded alphabet introducing 5-methylcytosine"

Supplementary Figure 1. The relationship between the value of N and the limit of corresponding information density for DNA data storage.

Supplementary Figure 2. Data recovery rate of six kinds of digital files under different error rates in repeated simulation tests.

Supplementary Figure 3. The relationship between the size of chosen file and attributes of oligos transcoded by R+

Supplementary Figure 4. Design flow of 8-nt adaptors.

Supplementary Figure 5. Gel electrophoresis image of sequence assembly in stages of group A and E.

Supplementary Figure 6. The proportion of substitutions, insertions and deletions for each position in oligo under the condition of no reference.

**Supplementary Tables**

Supplementary Table 1. The size of molecular alphabet and the corresponding limit of information density at various N values.

Supplementary Table 2. The concentration of final products of the step-by-step assembly of DNA sequences.

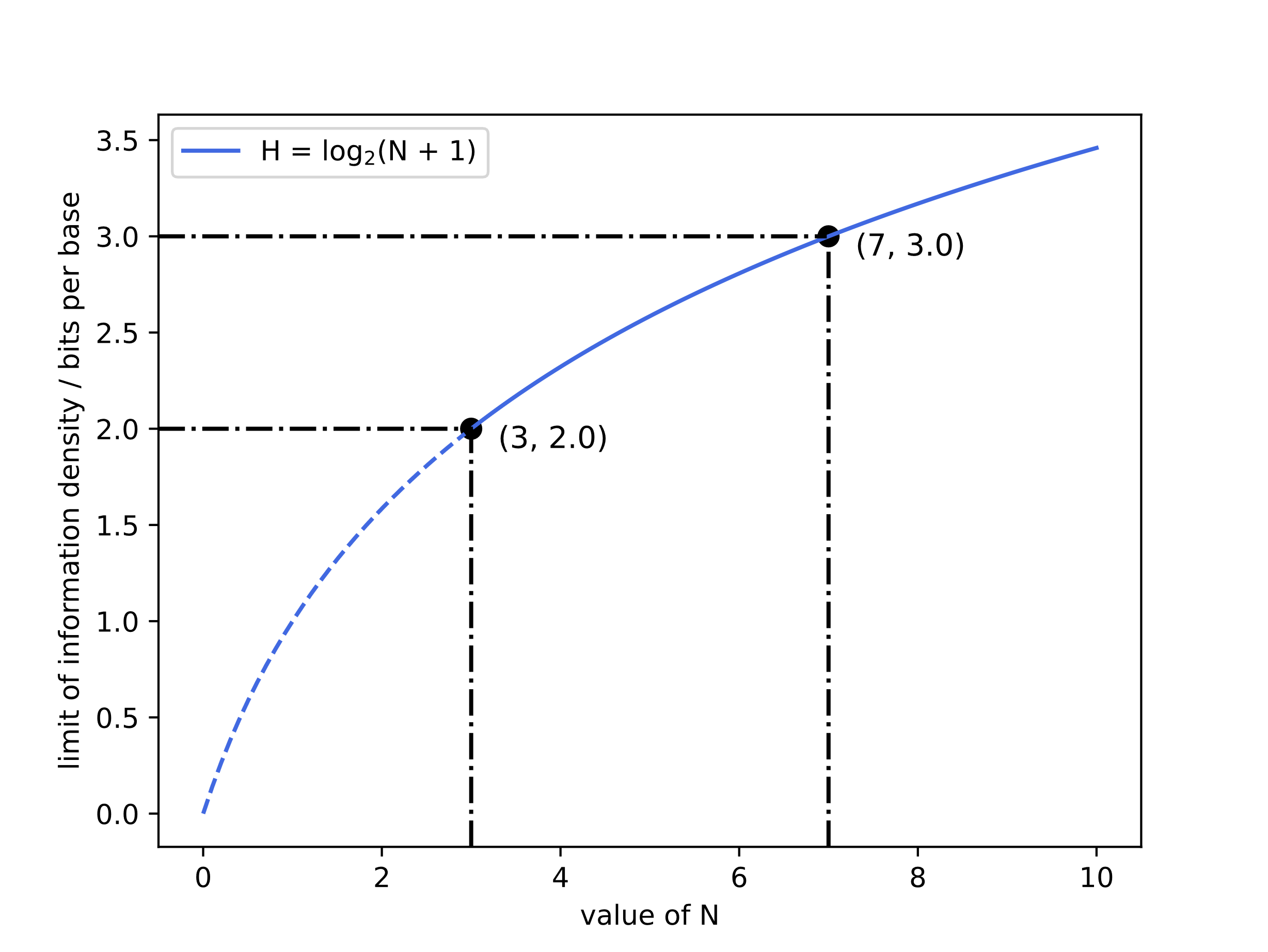

**Supplementary Figure 1. The relationship between the value of N and the limit of corresponding information density for DNA data storage.** From the information entropy formula, we could deduce that the information density limit of DNA data storage(H) has a logarithmic relationship with the size of molecular alphabet(N + 1). When the value of N is 3, the size of molecular alphabet is 4(i.e. A, C ,G and T), and the information density of the corresponding standard DNA data storage is limited to 2 bits/base. Similarly, the limit of information density is 3 bits/base when N is equal to 7.

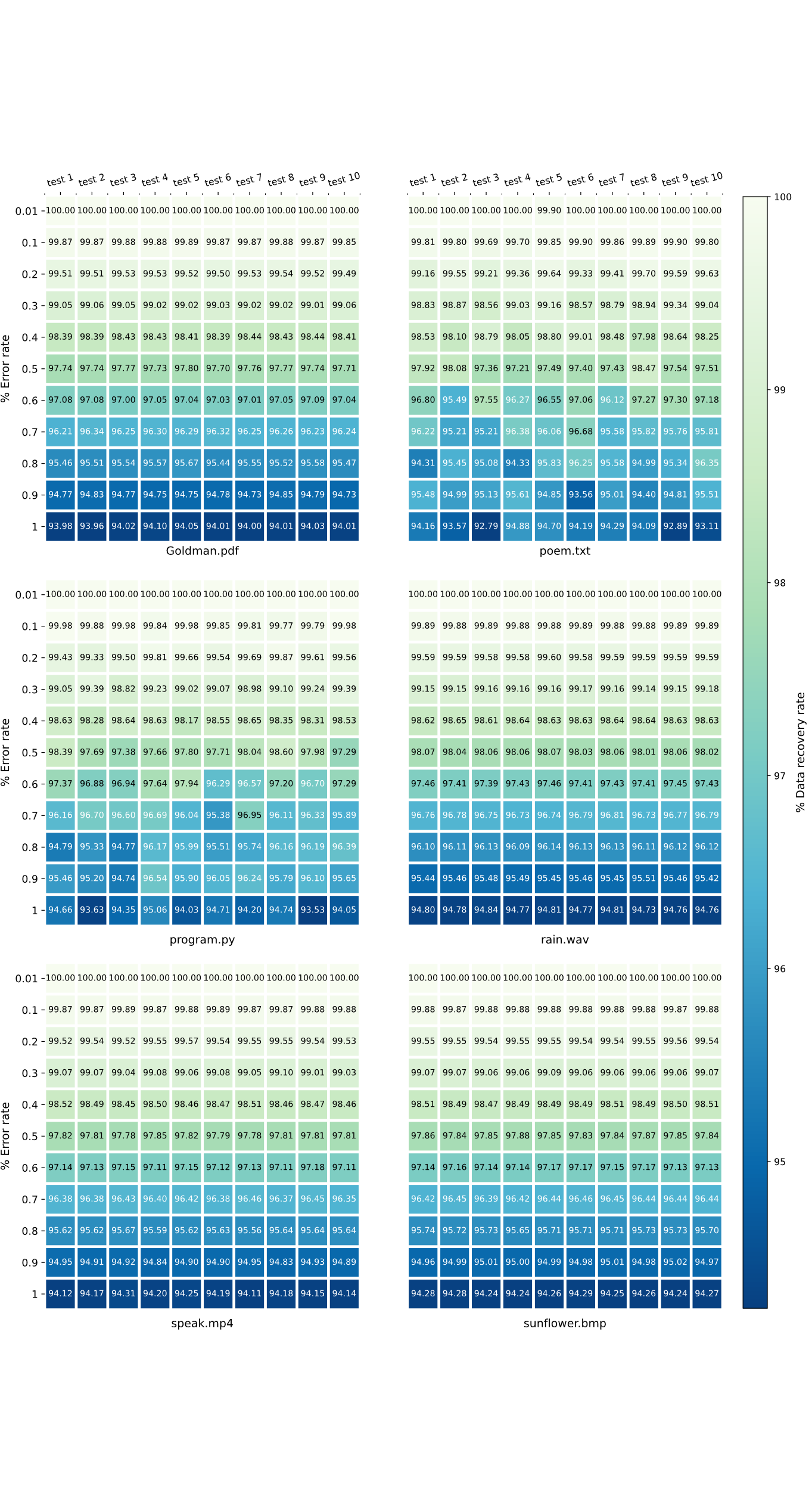

**Supplementary Figure 2. Data recovery rate of six kinds of digital files under different error rates in repeated simulation tests.** The data recovery rate for the six digital files is almost 100% at an error rate of 0.01%. With the increase of error rate, the data recovery rate in simulation test decreases gradually.

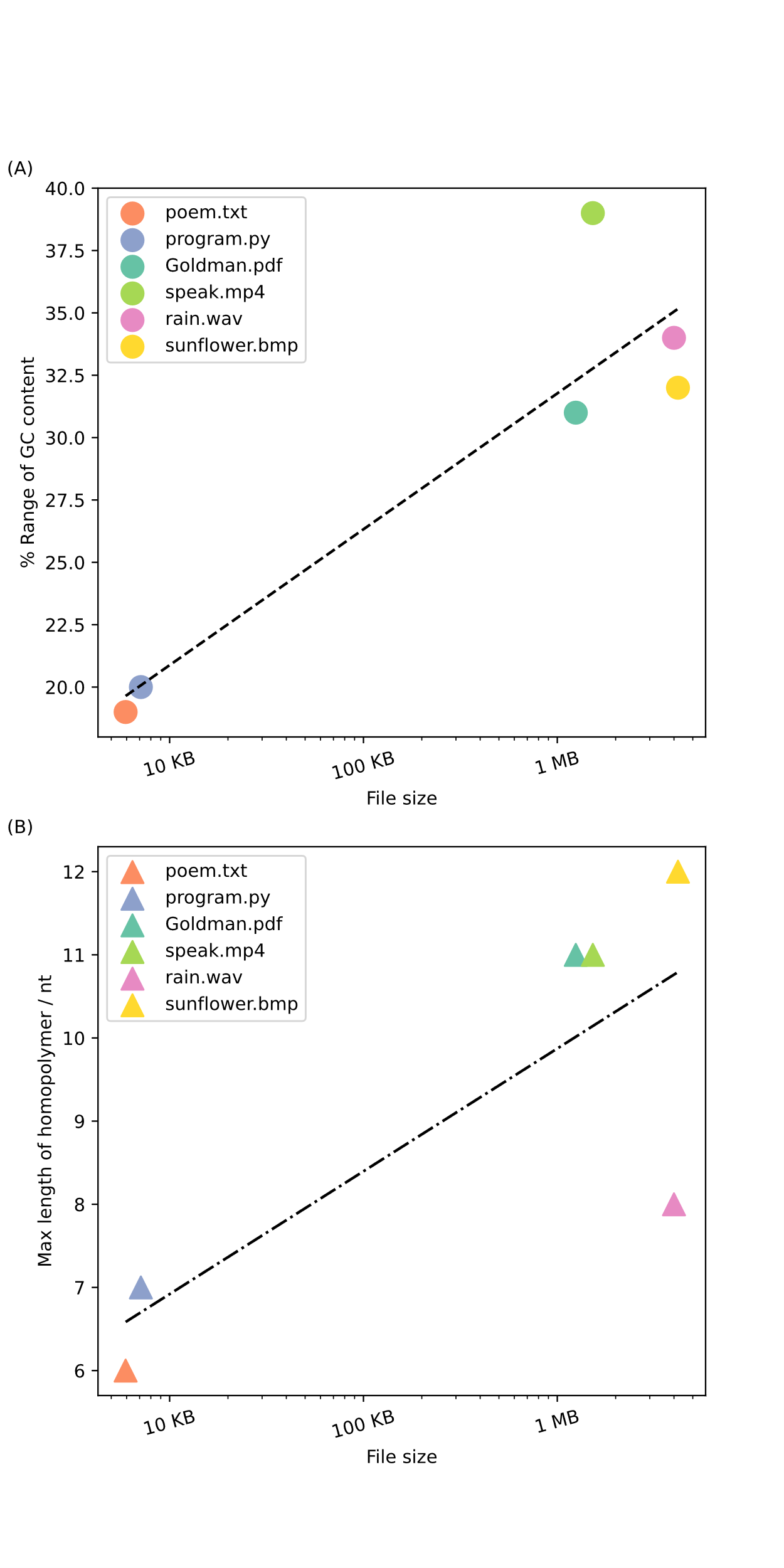

**Supplementary Figure 3. The relationship between the size of chosen file and attributes of oligos transcoded by R+.** (A) and (B) are respectively the relationship diagrams illustrating the growing tendency of GC content range and the maximum length of homopolymers as the file size increases.

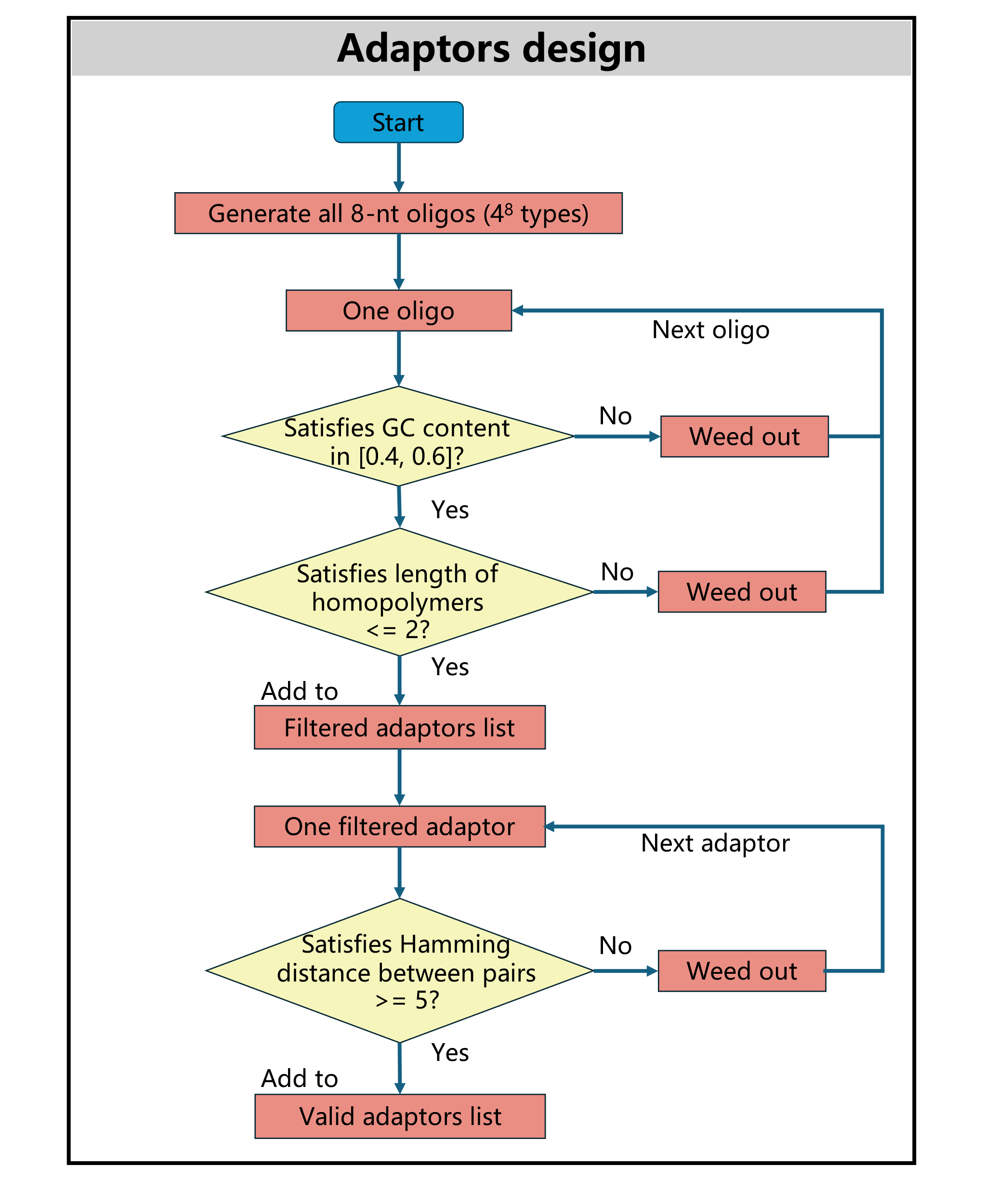

**Supplementary Figure 4. Design flowchart of 8-nt adaptors.** All 8-nt oligonucleotides as candidate sequences are subjected to a series of filtering criteria (GC content, length of homopolymers and Hamming distance) to select the valid adaptors, avoiding difficulties in biocompatibility and sequence ligation. Possible 8-nt oligos are generated firstly, totaling 4^8^ types. Next, the oligos undergo a filtering process where only those with a GC content between 40% and 60%, and the maximum homopolymer length not exceeding 2 nt, are retained as candidate adaptors. Finally, pairwise alignments are conducted among the candidate adaptors, and only those with a Hamming distance greater than 4 from all other candidates are valid ultimately.

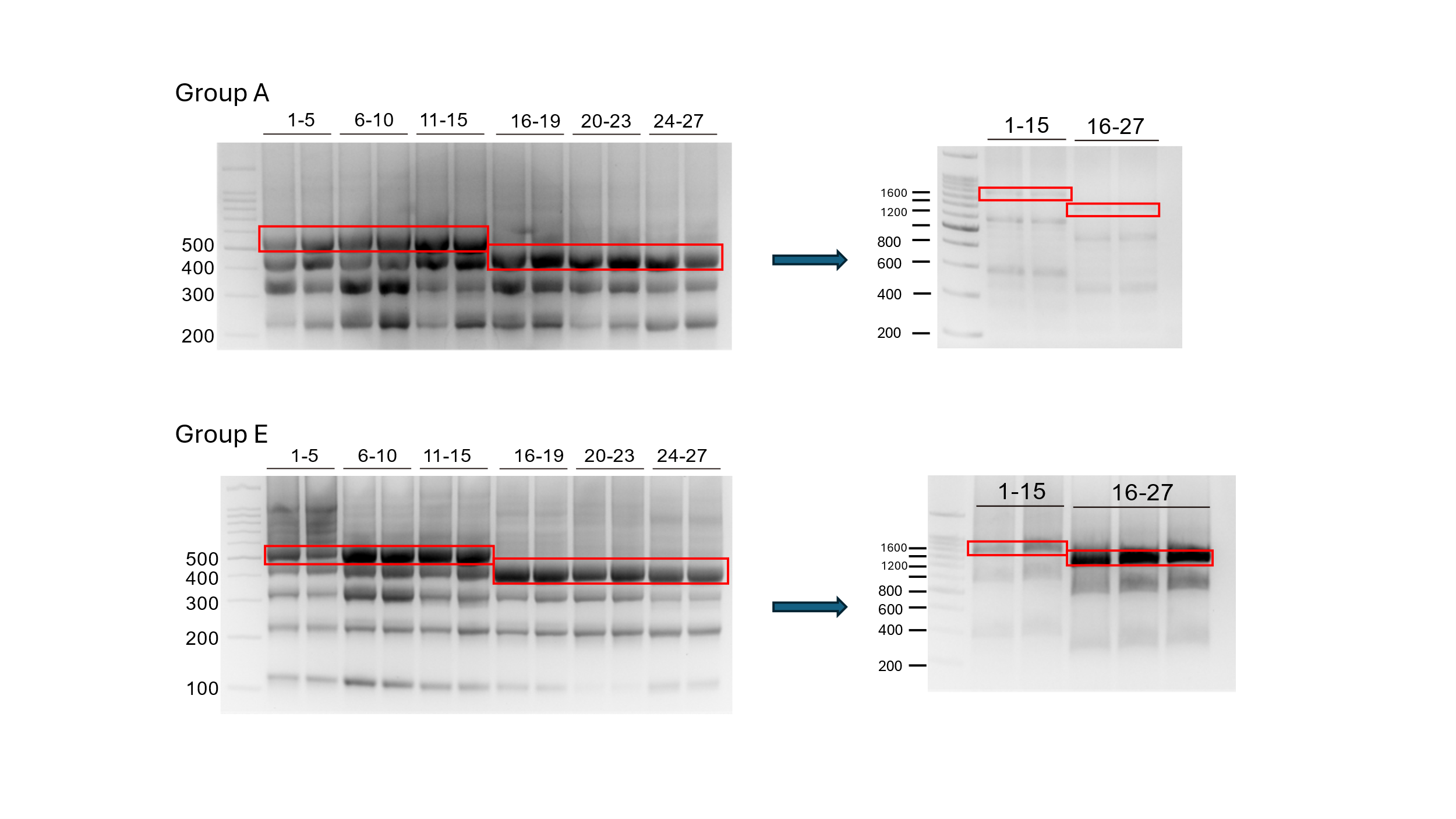

**Supplementary Figure 5. Gel electrophoresis image of sequence assembly in stages of group A and E.** The initial assembly of a group of oligos involves ligating every 5 of the first 15 oligos and every 4 of the remaining 12 oligos. The subsequent assembly ligates the first three and last three products obtained in the previous step.

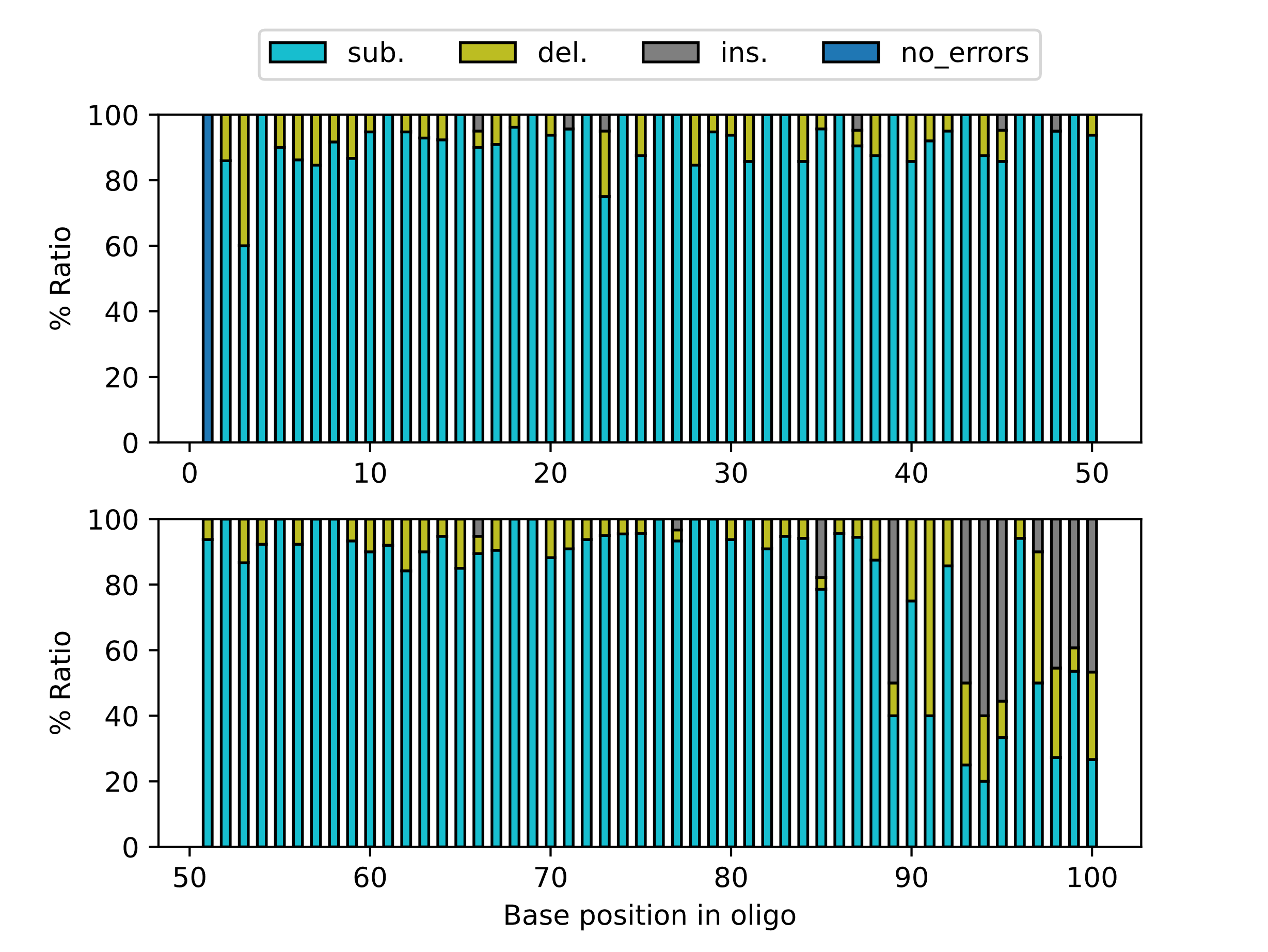

**Supplementary Figure 6. The proportion of substitutions, insertions and deletions for each position in oligo under the condition of no reference.**

**Supplementary Table 1. The size of molecular alphabet and the corresponding limit of information density at various N values.**

| Value of N | Size of molecular alphabet | Limit of corresponding information density for DNA data storage / bits per base |
| --- | --- | --- |
| 3 | 4 | 2.0 |
| 4 | 5 | 2.32 |
| 5 | 6 | 2.58 |
| 6 | 7 | 2.81 |
| 7 | 8 | 3.0 |
| N - 1 | N | log_2_N |

**Supplementary Table 2. The concentration of final products of the step-by-step assembly of DNA sequences.**

|  | **A** | **B** | **C** | **D** | **E** |
| --- | --- | --- | --- | --- | --- |
| Oligo 1-15 (ng/μl) | 6.14 | 28 | 6.02 | 5.52 | 15.5 |
| Oligo 16-27 (ng/μl) | 4.98 | 8.92 | 4.96 | 9.26 | 44 |
|  | **F** | **G** | **H** | **I** | **J** |
| Oligo 1-15 (ng/μl) | 14.8 | 17.3 | 17.2 | 6.98 | 13.6 |
| Oligo 16-27 (ng/μl) | 22.8 | 15.8 | 10.4 | 19.1 | 14.3 |
|  | **K** | **L** | **M** | **N** | **O** |
| Oligo 1-15 (ng/μl) | 16.3 | 21.8 | 10.7 | 9.2 | 26.8 |
| Oligo 16-27 (ng/μl) | 17.5 | 28.8 | 7.4 | 8.9 | 9.2 |
